# Dissociable developmental trajectories of Orbitofrontal subregion grey matter volume

**DOI:** 10.1101/2021.01.14.426730

**Authors:** S. G. Gibbons, M. P. Noonan

**Affiliations:** Department of Experimental Psychology, University of Oxford, Radcliffe Observatory, Anna Watts Building, Woodstock Rd, Oxford OX2 6GG; MRC Cognition and Brain Sciences Unit, University of Cambridge, 15 Chaucer Rd, Cambridge CB2 7EF

**Keywords:** Adolescent, Grey matter, Development, Voxel Based Morphometry, Human Connectome Project, Orbitofrontal Cortex

## Abstract

Adolescence is a period of development which is characterised by distinct differences in decision-making strategies relative to adults. While it is broadly established that there are relative differences in the structural maturation of the prefrontal cortex (PFC) and subcortical reward nuclei, such as the amygdala and ventral striatum, heterogeneity *within* the PFC is often neglected. In particular very little is known about the fine-scale gray matter (GM) development of the Orbitofrontal Cortex (OFC), itself critical to a number of learning and decision-making mechanisms which show delayed development trajectories. Here we applied voxel-based morphometry to examine subregional differences in OFC grey matter in high-quality structural MRI scans of 125 subjects aged 11-35yrs from the Human Connectome Project. First, we examined fine-scale GM maturation in 5 anatomically dissociable OFC subregions and identified the best-fitting polynomial model. Next, we directly compared developmental trajectories across 3 functionally dissociable subregions, revealing a complex topological developmental profile from medial to lateral subregions. Collectively, the two complementary analyses suggest that while unequivocally the phylogenetically younger lateral OFCs showed the greatest shift in GM volume across adolescence, with maturation continuing well into young adulthood, the differences between the medial and central OFC subregions suggested a more complex pattern of maturation than a simple graded medial to lateral topological development. We argue that knowledge of these fine-scale anatomical differences in maturation could explain precise mechanistic differences in goal-directed behaviours.

## Introduction

Adolescence is a late, sensitive period of development, during which young people gain independence and develop mature social goals (Fuhrmann *et al.*, 2015; van Duijvenvoorde *et al.*, 2016). Yet it is also a dangerous time of life, notoriously characterised by irrational decisions, that can often lead to accidental death (Viner *et al.*, 2012; Wolfe *et al.*, 2014), and coincides with the onset of many serious mental health illnesses (Paus *et al.*, 2008). Neuroanatomically, the teenage brain is also undergoing significant structural reorganisation (Paus *et al.*, 1999; Blakemore & Choudhury, 2006; Colby *et al.*, 2011; Whitaker *et al.*, 2016). The relative maturity of subcortical brain regions appears to overshadow the underdeveloped and poorly connected prefrontal cortex (PFC)(Mills *et al.*, 2014; Casey, 2015), potentially resulting in teenagers’ risky reward-seeking behaviour (Burnett *et al.*, 2010; Blakemore & Robbins, 2012; Galvan & McGlennen, 2013). While the importance of the well-established finding of the late maturation of the PFC cannot be understated, the PFC is composed of a number of fundamentally distinct sub-regions (Passingham & Wise, 2012; Sallet *et al.*, 2013; Neubert *et al.*, 2014; Neubert *et al.*, 2015b) and this heterogeneity is neglected in current whole-brain developmental maps of cortical maturation (Sowell *et al.*, 2003; Gogtay *et al.*, 2004). As we move to a circuit-based understanding of learning and decision-making (Hunt & Hayden, 2017) and the altered network states during adolescence (Casey, 2015), a critical fundamental advancement then becomes the fine-scale quantification of the developmental of anatomically dissociable PFC subregions.

One particularly developmentally delayed subregion of the PFC is the orbitofrontal cortex (OFC), a region critically involved in learning and decision-making, which broadly does not reach gross maturity until the early 20s (Gogtay *et al.*, 2004). The OFC is itself comprised of numerous cytoarchitectonically and connectionally distinct subregions (Carmichael & Price, 1996; Neubert *et al.*, 2015b). Using diffusion weighted imaging, Neubert and colleagues created detailed parcellation maps based on the unique connectivity profiles of each subregion and delineated 21 key human orbital and medial frontal brain regions, of which five anatomically dissociable regions are consistently considered part of the orbitofrontal cortex itself. The first corresponds to medial area 14 which extends between the medial orbitofrontal sulcus and the rostral sulcus in ventromedial prefrontal cortex, as such it is often referred to as mOFC/vmPFC in humans (Ongur & Price, 2000). Laying ventral to area 14, is area 11m. Area 11m encompasses the gyrus rectus and cortex beneath, and medial to the olfactory tract. Despite area 11m laying medial to area 13 and extending rostrally onto the medial surface, its projections are more interlinked with the more central and lateral orbital networks (Carmichael & Price, 1996) and as such it is not considered part of mOFC/vmPFC. Area 11 (excluding area 11m), which lies immediately posterior to the frontopolar region of area 10, and the more posterior area 13 are located laterally from the medial orbitofrontal sulcus and used to be referred to as the lateral orbitofrontal sulcus are now referred to as central OFC (Rudebeck *et al.*, 2017; Sallet *et al.*, 2020). Central OFC is now also distinguished from cortex beyond the lateral orbitofrontal sulcus which transitions into ventrolateral prefrontal cortex and aligns mostly with the gyral region of the orbital part of inferior frontal gyrus. This corresponds to the orbital part of area 12 (12o) in macaques and Brodmann’s area 47o in humans (referred to from here as area 47/12o).

The anatomically-defined dissociations between OFC subregions are complemented by evidence of fine-scale functional dissociations. For example, lesion studies support in adult humans and macaques show distinct functional contributions of medial area 14, central OFC (areas 11 and 13, typically lesioned together) and lateral OFC area 47/12o in learning and decision-making tasks. These studies demonstrate that mOFC/vmPFC in both species is necessary for decisions that involve multiple valuable options to choose between and for focusing a decision on the most relevant information (Noonan *et al.*, 2010; Vaidya & Fellows, 2016; Noonan *et al.*, 2017). Central OFC regions are involved in the updating of object valuations according to current motivational states (Rudebeck *et al.*, 2013b). Finally, the lateral regions of the OFC are critical for behavioural flexibility and learning the consequences of choices (Noonan *et al.*, 2010; Noonan *et al.*, 2011; Rudebeck *et al.*, 2013b; Noonan *et al.*, 2017; Sallet *et al.*, 2020). To date, fine-scale developmental differences in subregion maturation have not been investigated but quantifying these differences is likely to be key in interpreting the behavioural developments in contingent learning (Javadi *et al.*, 2014; Hauser *et al.*, 2015) and value-guided decision-making (Kwak *et al.*, 2015), as well as more broadly advancing our understanding of the maturation of reward circuits in adolescents (Galvan *et al.*, 2005). Therefore the current study combined recently released high-quality HCP developmental data (Somerville *et al.*, 2018) with the HCP young adult dataset (Van Essen *et al.*, 2013), and used voxel based morphometry methods (Good *et al.*, 2001) to identify and quantify relative grey matter (GM) differences in fine-scale OFC subregions across an age range of 11-35yrs.

## Methods

### Participants

125 individuals were sampled from the HCPYA (*n* = 56) and HCPD (*n* = 69) databases. Participants were aged 11-35 years and categorised into three age groups, 11-17, 18-23 and 24-35 years old. These categories were informed by societal and neuroanatomical developmental changes (Viner *et al.*, 2012; van Duijvenvoorde *et al.*, 2016) with particular acknowledgement of the late maturation of the OFC (Gogtay *et al.*, 2004). Approximately equally-sized populations were sampled within these age ranges in order to create a non-biased study specific GM template. 43 subjects, aged 11-17 years from HCPD had pre-processed T1w images available at the time of analysis. We matched this sample size for the 24-35 age group, randomly selecting 43 subjects with T1w images and necessary demographic data from HCPYA. Finally, bridging upper and lower ranges of HCPD and HCPYA, we populated the 18-23 age group by randomly sampling 26 HCPD subjects between 18-21 years and 13 HCPYA subjects between 22-23 years. The non-biased study-specific template was generated by randomly selecting the N of the smallest group (*n* = 39) from each of the larger groups (see Figure 1 for a histogram of participant ages). Demographic data, sex and handedness (overall score of number of responses of right hand minus those of left hand, divided by right plus left, using items of the Edinburgh handedness inventory (Oldfield, 1971)) for all participants can be found in Table 1.

**Figure 1:**
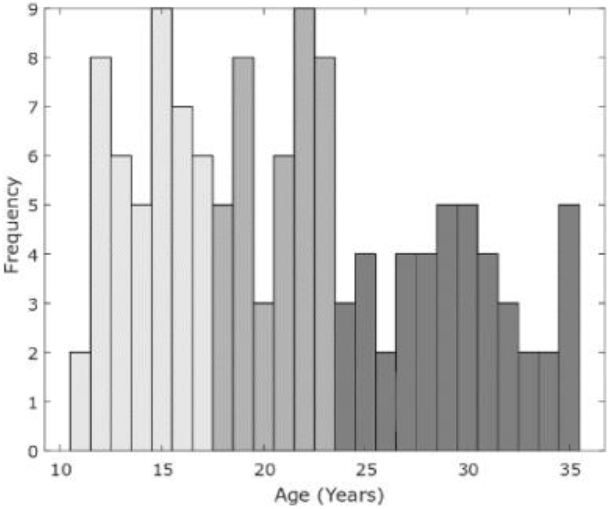
Histogram of age of participants in voxel-based morphometry analysis. For the purposes of template creation, participants were categorised a priori into three age groups: 11-17 (light grey); 18-23 (mid grey); 24-35 (dark grey).

**Table 1:** Participant demographic data.

|  | *All* | *11-17 years* | *18-23 years* | *24-35 years* |
| --- | --- | --- | --- | --- |
| *Total N* | 125 | 43 | 39 | 43 |
| *Sex (N Female/Male)* | 67/58 | 21/22 | 18/21 | 28/15 |
| *Handedness (N right/left)* | 115/10 | 38/5 | 37/2 | 40/3 |

### Image Acquisition and Minimal Pre-Processing

T1w structural images were acquired with HCP acquisition parameters (Glasser et al., 2013): 256 sagittal slices were acquired with TR = 2400ms, TE = 2.14ms, flip angle = 8°, FOV = 224mm, matrix = 320. The primary aim of these selected parameters was to achieve high spatial resolution: T1w images had a voxel size of 0.7mm isotropic. All structural images underwent minimal pre-processing via the three HCP structural pipelines *PreFreeSurfer*, *FreeSurfer*, and *PostFreeSurfer*. The PreFreeSurfer pipeline generates a *Native Volume Space* brain-extracted, undistorted T1w image which is linearly aligned to the axes of MNI space. Full details of structural image acquisition and minimal pre-processing pipelines are reported by Glasser et al (2013).

### Voxel Based Morphometry

Based on the voxel-based morphometry (VBM)(Douaud *et al.*, 2007) analysis pipeline, we conducted analysis to identify fine-scale differences in GM between OFC subregions across adolescence and early adulthood. The HCP-provided skull stripped T1w images were individually segmented into GM, white matter (WM) and cerebral spinal fluid (CSF) before being affine-registered to the GM ICBM-152 template using FLIRT (Jenkinson & Smith, 2001) followed by nonlinear registration using FMRIB’s Nonlinear Image Registration Tool (FNIRT)(Andersson *et al.*, 2008). Of the resulting images, those randomly selected to compose the non-bias study-specific template (equal number of subjects from each of the age ranges 11-17, 18-23 and 24-35 years) were averaged, the native grey matter images of all subjects were then non-linearly re-registered to this template, and concatenated into a 4D image. The resulting unsmoothed 4D image is modulated for the contraction or enlargement of each subject’s image to the template with each voxel of each image being multiplied by the Jacobian of the warp field (Good *et al.*, 2001). The final step of our VBM analysis was to extract all subject’s values from the 4D GM modulation image – from here on these values will be referred to as “GMm” - for each of our ROIs, providing a between-subjects comparison of volume which could be regressed against age to map developmental trajectory. ROI average Jacobian values were extracted using fslmeants in combination with the corresponding mask. Higher GMm values correspond to more GM, reflecting regional expansion of the native GM images relative to the template, whereas lower values correspond to less GM and reflect regional GM contraction of native images.

### Regions of Interest

OFC subregions were selected a-priori on the basis of Neubert et al.’s (2015a) parcellation of the human medial and orbital regions of the prefrontal cortex. Parcellations were identified according to functional connectivity fingerprinting, with Neubert and colleagues identifying five anatomically dissociable OFC subregions: Brodmann’s areas 11, 11m, 13, 14, and 47/12o (Figure 2). As introduced above, the OFC is also often dissociated based on functionality derived from similarities in structural connectivity (Carmichael & Price, 1996) or findings from lesion studies (Noonan *et al.*, 2010; Noonan *et al.*, 2017; Rudebeck *et al.*, 2017). Consequently, we grouped two of these subregions into central [11 & 13] OFC and compared GM maturation with the medial OFC [14] and lateral OFC [47/12o] (Figure 2).

**Figure 2:**
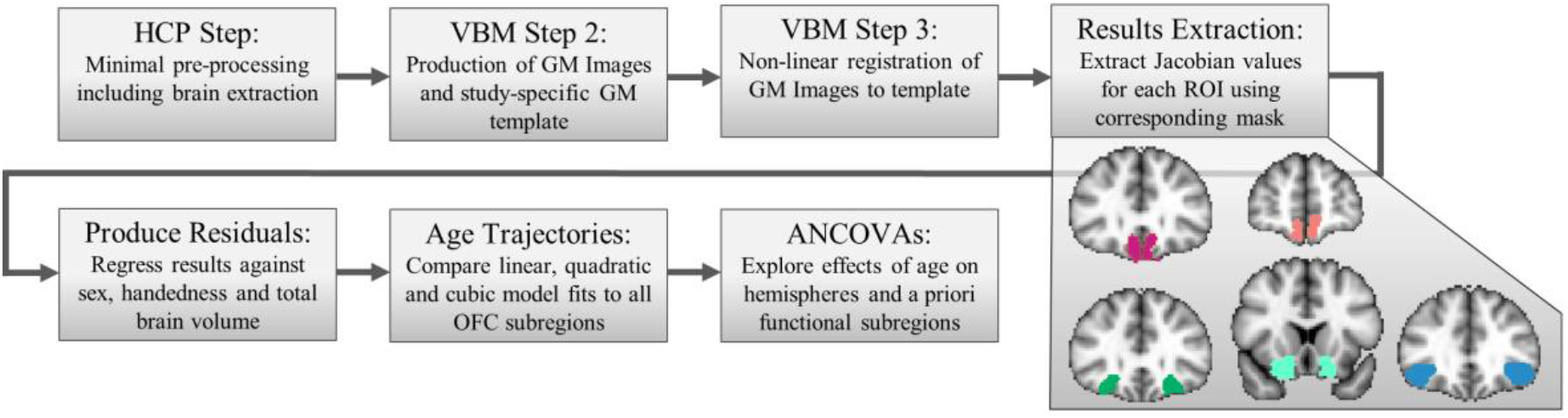
Workflow of methodology and the five anatomical Orbitofrontal Cortex parcels (top left to bottom right: 14, 11m, 11, 13, 47/12o) from (Neubert et al., 2015b). Abbreviations: HCP, Human Connectome Project; VBM, Voxel Based Morphometry; GM, grey matter; ROI, region of interest; OFC, Orbitofrontal Cortex.

### Statistical Analysis

#### Partialing out confounding covariates

To control for the effects of subjects’ sex, handedness and total brain volume, these variables were regressed out of the GMm values for each of the five parcels across each hemisphere using MATLAB (2014a, The Mathworks, Inc., Natick, MA, United States). The resulting residual GMm were used in analyses.

#### Correlations and Model Fits

We conducted bivariate correlations to investigate the relationship between age and GMm of each of the five anatomically defined OFC subregions in each hemisphere. After demonstrating that such a correlation exists for the specific subregion, we then fitted linear, quadratic and cubic functions, using lowest Akaike Information Criterion (AIC) value to identify the best-fitting model, and adjusted R^2^ to metricise goodness-of-fit. In addition, we ran a likelihood-ratio test to confirm whether the best fitting model explained significantly more variance than a null model. The null model was constructed by assuming no relationship with age and instead fitting the mean of the GMm.

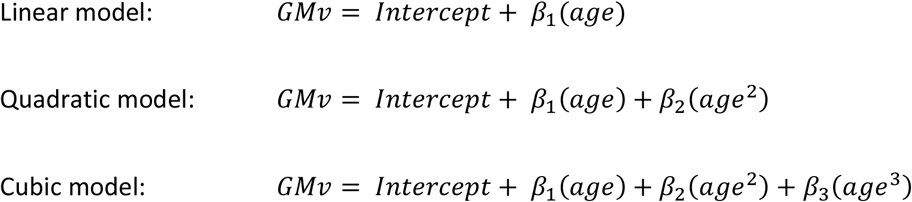

Finally, we used the best-fitting model estimate of the residual GMm values at ages 11yrs and 35yrs to identify the percentage change in grey matter volume (GMv) for each functional subregion in the age range sampled using the following function: (model estimate of GMm age 11yrs - model estimate of GMm age 35yrs)*100. These follow-up analyses allowed us to characterise differences in shape and steepness between subregions.

#### Analysis of Covariance

To investigate and compare GM trajectories across functionally dissociable subregions of the OFC we subjected medial area 14, central OFC (cOFC average GMm of area 11 and area 13) and lateral area 47/12o to a repeated measures ANCOVAs. All key assumptions necessary to subject the data to a repeated measures ANCOVA were tested and met. With both the dependent variable, GMm, and the covariate, age, measured on continuous scales, and the independent variables being subregions of OFC and hemispheres. No significant outliers were identified (all data were less than 3 standard deviations from the mean, except two subjects with 2 data points <4SD above the mean, and two subjects with 1 data point <4×SD above the mean). Residuals were approximately normally distributed for each category of the independent variable (Shapiro-Wilk tests p > 0.05), with the exception of area right hemisphere 14m (Shapiro-Wilk test p = 0.019). The covariate was linearly related to the dependent variable at each level of the independent variable (ps ≤ 0.045), with the exception of GMm values from the right cental OFC (p = 0.808). Homoscedasticity was confirmed via scatterplots of the standardized residuals against the predicted values and all coefficient VIF collinearity statistics reported as <10.

Residuals were subjected to ANCOVA methodology outlined by Schneider and colleagues (2015). The covariate measure of age was manually centered across all of the participants in the experiment before entered into 2-way repeated measures ANCOVAs. We included OFC subregion and brain hemisphere as independent variables, with age as a covariate (3(ROI: cOFC[11 & 13], area 14 and area 47/12o) × 2(Hemisphere) × age ANCOVA), and examined evidence in favour of the hypothesis of a differential effect of age on OFC subregion GMm, i.e. an interaction between OFC subregion GMm and age. Schneider et al (2015) then advise employing a standard ANOVA (3(OFC subregion) × 2(hemisphere) to evaluate any effects not involving the age covariate. However, due to the study-specific GM template being an age-wise representative average of the brain structure of the subjects studied, and GMm values being relative to this average template (zero), the mean of all values will always be approximately zero for each of the OFC subregions. We therefore did not anticipate any independent main effects of OFC subregion or hemisphere. In this analysis, the primary focus examines *relative* differences in GMv trajectories and percentage changes in GMv, not differences in absolute GMv between brain regions. Accordingly, we do not report the results of follow up ANOVA.

We next examined the three functionally dissociable subregions in three separate 2(ROI: cOFC vs area 14 | cOFC vs area 47/12o | area 14 vs area 47/12o) × 2(Hemisphere) × age ANCOVAs. Again, we do not report results of follow uppost-hoc ANOVAs.

## Results

### GM developmental model fits for all anatomical OFC subregions

To investigate the relationship between age and GM of each of the five anatomically defined OFC subregions we performed bivariate correlation analyses with complementary model fit comparisons. While GMm in left and right hemispheres of each OFC subregion were significantly positively correlated (all *r*s > 0.532, *p*s < 0.001) we fit models to the GMm separately for each hemisphere of all five OFC subregions. Figure 3 illustrates the GMm values for each OFC subregions correlated against age and, if this correlation is significant, fit with their respective best fitting polynomial model.

**Figure 3.**
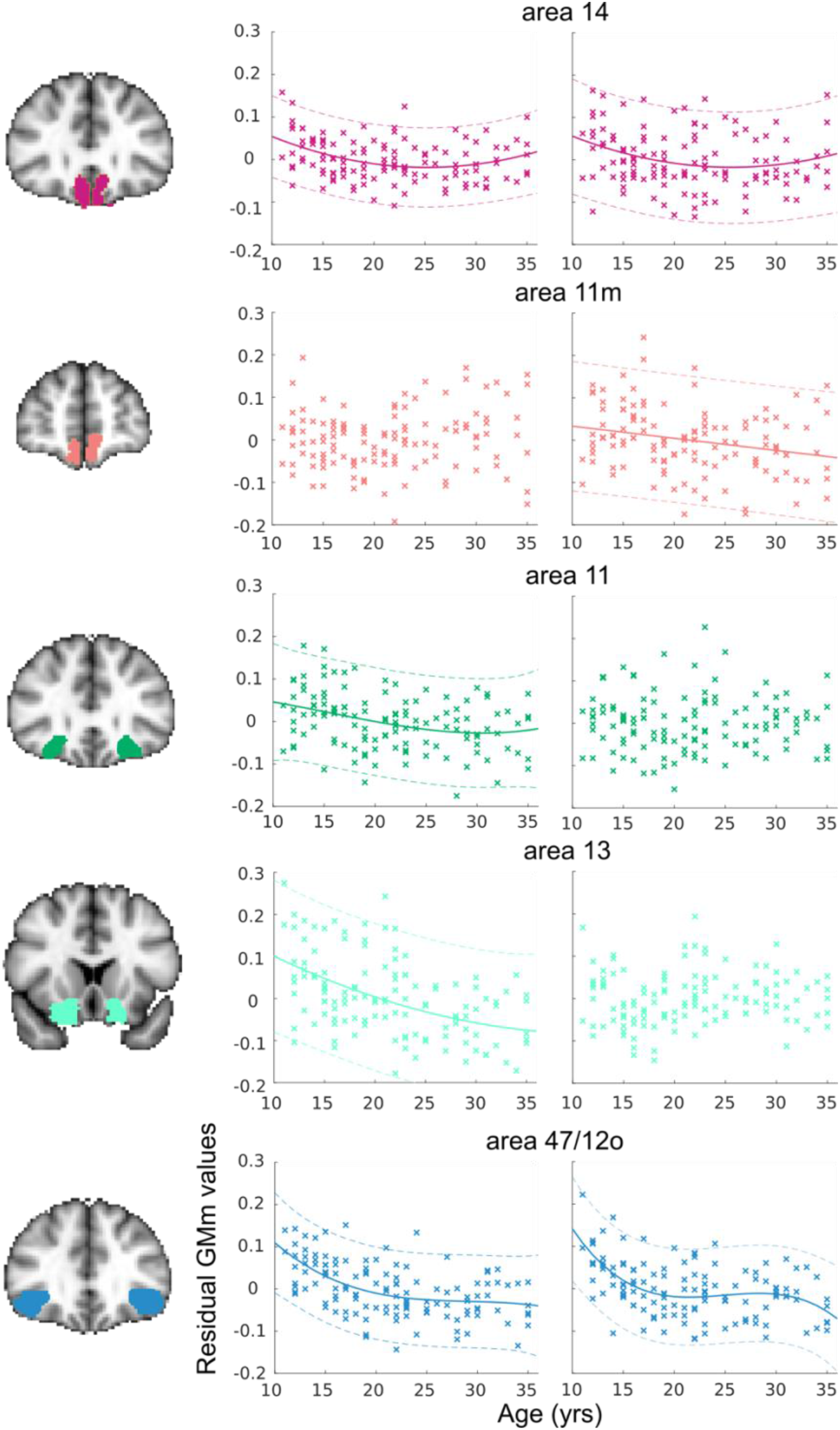
OFC subregion masks from Neubert and colleagues (2015b) used to extract GMm values for each subregion following the voxel-based morphometry analysis. GMm values for each OFC subregion, accounting for sex and handedness were correlated against age. Left and right hemispheres are shown separately in the respective panels. Results indicate a heterogenous pattern of development across the five OFC subregions and between hemispheres. Where there is a significant correlation with age, data is fit with the best fitting polynomial model (solid line) and 95% confidence intervals (dashed lines). Individual participant data points are shown as ‘x’s.

**Figure 4.**
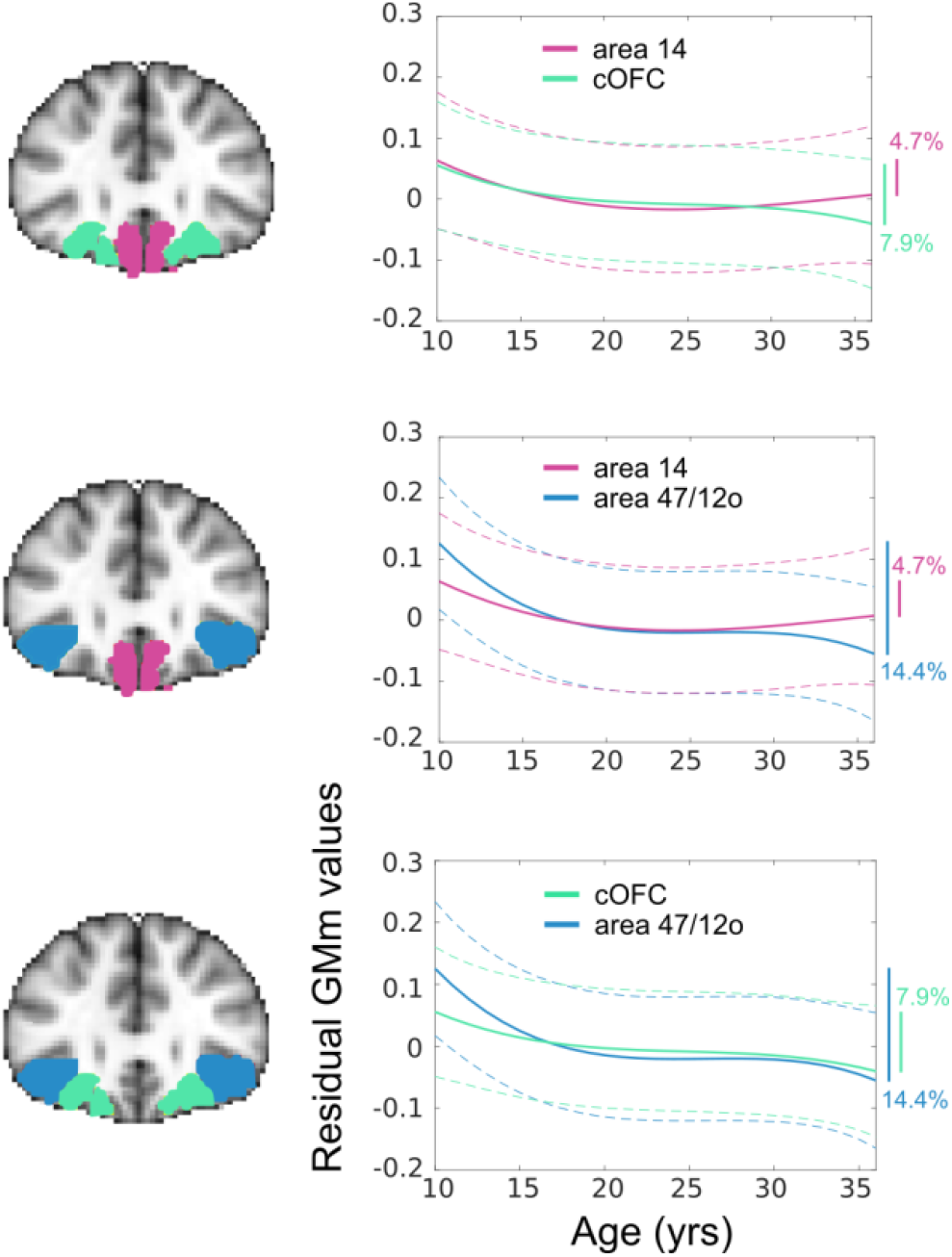
OFC subregion masks from Neubert and colleagues (2015b) for the pair-wise comparisons of the residual GMm values extracted from the three functionally dissociable OFC subregions; medial area 14, central OFC (cOFC; area 11+13) and lateral area 47/12o. Best model fits (solid line) and 95% confidence intervals (dashed lines) are shown for each OFC subregion against each other. Model fits were used to estimate the percentage difference in GM volume between the youngest and oldest participants, reported on the right of each subplot. Lines to the right represent the proportionate reductions in GM volume from the youngest to the oldest age sampled. The pattern of results suggests a loose graded maturation of GM from medial to lateral areas as indexed by the differences between subregion developmental trajectories, with the clearest difference between the most medial OFC subregion (area 14) and the most laterally considered region (area 47/12o).

Medial area 14 showed a significant relationship between GMm and age in both hemispheres (area 14l: *r*_124_ = −0.22, *p* = 0.02; area 14r: *r*_124_ = −0.18, *p* = 0.05), with AIC determining these were best characterised by quadratic functions (area 14l: adjusted *R^2^*= 0.13, LR *p* < 0.001; area 14r: adjusted *R^2^* = 0.07, LR *p* = 0.004). By contrast, only the right hemisphere of OFC subregion 11m correlated significantly with age (area 11mr: *r*_124_ = −0.25, *p* < 0.001) and was best fit with a linear model, though according to the likelihood ratio, not better than the null model (area 11mr: adjusted *R^2^*= 0.06, LR *p* = 1.00). In central OFC subregions, GMm in the left hemispheres significantly negatively correlated with age (area 11l: *r*_124_ = −0.32, *p* < 0.001, area 13l: *r*_124_ = −0.48, *p* < 0.001), with AIC suggesting they were best characterised by a cubic function (area 11l adjusted *R^2^*= 0.10, LR *p* < 0.001), and a quadratic function (area 13l: adjusted *R^2^*= 0.23, LR *p* = 0.044) respectively. GMm in the right hemispheres of both central OFC subregions was not correlated with age. Finally, lateral area 47/12o GMm also negatively correlated with age in both hemispheres (area 47/12ol: *r*_124_ = −0.48, *p* < 0.001; area 47/12or: *r*_124_ = −0.40, *p* < 0.001) with the trajectory best characterised by a cubic function in both hemispheres (area 47/12ol: adjusted *R^2^*= 0.28, *p* < 0.001; area 47/12or: adjusted *R^2^*= 0.27, *p* < 0.001).

### Quantifying and comparing GMm across a priori functional OFC subregions

Next, we sought to test the key hypothesis that three functionally distinguishable OFC subregions mature at different rates. First, we quantified differences in grey matter between the youngest and oldest sampled participants in medial area 14, central OFC (average GMm of area 11 and area 13), and lateral area 47/12o. Averaging GMm across hemispheres and using the same model fit procedure describe above, we showed that medial, central and lateral regions were best fit, according to the AIC, each by a cubic model (area 14: adjusted R^2^= 0.097, LR *p* < 0.001, cOFC: adjusted R^2^= 0.107, LR *p* < 0.001, 47/12o: adjusted R^2^= 0.310, LR *p* < 0.001). Taking these model fits we determined the difference in GM across the age range sampled and report that area14 has a 4.69% reduction while cOFC shows a 7.86% reduction in GMv from age 11yrs to 35yrs. Note, when the relationship between the left cOFC is assessed in isolation this subregion was also best fit by a cubic model (adjusted R^2^= 0.279 LR *p* < 0.001) and showed a 10.91% reduction in GMv. By far the greatest reduction in GMv is in lateral area 47/12o which showed a 14.44% loss across the age range tested.

To examine whether these differences in GMv between subregions were meaningful we conducted ANCOVA analyses, comparing differences in GMm as a function of age among the three a priori functional OFC subregions of interest. GMm values from medial area 14, cOFC (area 11 and area 13) and lateral area 47/12o from both hemispheres were subjected to a 3 (OFC subregion) × 2 (hemisphere) × age repeated measures ANCOVA. The results support the primary hypothesis that GMm across the three different functionally dissociable subregions is differentially affected by age (subregion × age interaction: *F*_2,246_ = 5.608, *p* = 0.004, *η^2^*= 0.044, Power = 0.855) and the two factors differentially interact with hemisphere (subregion × hemisphere × age interaction: F_2,246_ = 20.354, *p* < 0.001, *η^2^*= 0.142, Power = 1.000). Age itself is also a significant explanatory variable of GMm overall (main effect of age: *F*_1,123_ = 29.415, *p* < 0.001, *η^2^*= 0.193, Power = 1.000) and interacts with hemisphere independently (*F*_1,123_ = 24.160, *p* < 0.001, *η^2^*= 0.164, Power = 0.998).

We performed follow up comparisons of each functional subregion with each other and report the key effects below.

#### Medial area 14 v Lateral area 47/12o

We compared area 14 and 47/12o subregions directly using ANCOVA analysis. The results reveal differential developmental trajectories of GMm between these subregions as indexed by a significant age × OFC subregion interaction (F_1,123_ = 9.896, p = 0.002, η^2^= 0.074, Power = 0.877). Age was also a significant independent predictor of GMm (F_1,123_ = 26.437, p < 0.001, η^2^= 0.177, Power = 0.999). There were no further interactions involving age (age × hemisphere interaction: F_1,123_ = 0.239, p = 0.626, η^2^= 0.002, Power = 0.077; age × OFC subregion × hemisphere interaction: F_1,123_ = 0.768, p = 0.382, η^2^= 0.006, Power = 0.140). Combined with the results of model fitting and calculation of percentage GMv change, these results suggest lateral area 47/12o has relatively more GMv change than medial area 14 during adolescence, showing greater GMv development across the age-range sampled, and a statistically dissociable developmental course.

#### Medial area 14 v Central OFC

Comparing GMm between medial area 14 and cOFC using ANCOVA analysis revealed that despite a significant main effect of age (main effect of age: F_1,123_ = 13.191, *p* < 0.001, *η^2^*= 0.097, Power = 0.950) there was not a differential GMv modulation with age across the two subregions (age × OFC subregion interaction: F_1,123_ = 1.183, *p* = 0.279). However, the interaction effect between hemisphere and age (age × hemisphere interaction: F_1,123_ = 29.949, *p* < 0.001, *η^2^*= 0.196, Power = 1.000) and the 3-way interaction with OFC subregion were significant (age × OFC subregion × hemisphere interaction: F_1,123_= 28.336, *p* < 0.001, *η^2^*= 0.187, Power = 1.000). Post hoc ANCOVAs revealed significant differences in GMm between area 14 and cOFC in the right hemisphere (F_1,123_ = 4.370, p = 0.039, *η^2^*= 0.034, Power = 0.545), but substantially larger developmental differences in left hemispheres (F_1,123_ = 23.490, p < 0.001, *η^2^*= 0.160, Power = 0.998). This pattern of effects may underlie the pattern of interactions seen in the omnibus ANCOVA. Combining these results with the model fits and GMv percentage change analysis, it is apparent cOFC undergoes a complex pattern of development. Left cOFC appears to experience greater GM pruning than the left hemisphere of medial area 14 across the age range studied, while right cOFC shows no relationship with age, potentially indicating maturation has reached a more stable phase.

#### Lateral 47/12o v Central OFC

Finally, ANCOVA analysis revealed a complex relationship between lateral area 47/12o and cOFC developmental trajectories. In addition to age being a significant predictor of GMm in both subregions (main effect of age: F_1,123_ = 38.873, *p* < 0.001, *η^2^* = 0.240, Power = 1.000), there was also a significant interaction between OFC subregions and age (subregion × age: F_1,123_ = 4.753, p = 0.031, *η^2^*= 0.037, Power = 0.581). There was also a significant interaction between hemisphere and age (age × hemisphere interaction: F_1,123_ = 36.764, *p* < 0.001, *η^2^*= 0.230, Power = 1.000) and a further interaction with OFC subregion (age × OFC subregion × hemisphere interaction: F_1,123_ = 27.162, *p* < 0.001, *η^2^*= 0.181, Power = 0.999). Post hoc ANCOVAs showed that this interaction was due to significant differences in trajectories between the two subregions in the right hemisphere (F_1,123_ = 19.37, p < 0.001, *η^2^*= 0.136, Power = 0.992) but not the left hemisphere (F_1,123_ = 0.36, p = 0.552). While difficult to fully interpret without sampling over a wider range of participant ages, this effect might suggest GMv in the right hemisphere of the cOFC has already matured by approximately 11 years (as there is no a relationship with age), driving the significant differences with right hemisphere of area 47/12o, but that the left hemisphere cOFC by contrast was still maturing. In the left hemisphere, GM maturation of the cOFC and area 47/12o do not differ in terms of the overall timing.

## Discussion

The aim of this study was to quantify the relative differential patterns of structural maturation between anatomically and functionally dissociable OFC subregions. Analyses of high-quality structural MRI scans of 125 Human Connectome Project subjects of adolescent and young adult brains shows unique developmental profiles for each anatomically dissociable subregion. While there are fine-scale regional and hemispheric differences in the maturation profiles within the orbitofrontal cortex, the pattern of results provide evidence of a loose graded maturation with the phylogenetically younger lateral area 47/12o showing a longer developmental maturation profile compared medial OFC and showing a significant decrease in GMv across adolescence that continues well into young adulthood.

At the more fine-scale level, we observed that medial OFC showed a relatively small percentage reduction in GMv across age compared to other functional regions and the relationship with age was best characterised by quadratic functions. Central OFC subregions 11 and 13, though phylogenetically younger than their medial neighbour, showed hemispherically heterogeneous relationship between GMv and age. Their relationship with age was best fit by cubic and quadratic functions respectively, when it showed a significant relationship with age at all. The more lateral region 47/12o, however, unequivocally showed the greatest percentage reduction in GMv, with GM maturation in both hemispheres best characterised by cubic functions indicating protracted development. Comparing the developmental trajectories across functionally heterogenous subregions, it is apparent that the maturation of the area 47/12o was consistently dissociable from that of medial OFC. Medial OFC is also consistently dissociable from the cOFC but for different reasons in each hemisphere: with the right hemisphere cOFC potentially having reached a plateaux of GM maturation, while the left is still developing. Finally, area 47/12o is only dissociable from right cOFC, but not the still maturing left cOFC. At this level, the pattern of results speaks against a categorical maturation of GM between functionally dissociable OFC subregions and instead add a layer of complexity to a simple graded medial to lateral GM maturation.

Such differences in subregion developmental trajectory could explain developmental differences in the influence of the distinct decision-making mechanisms that these subregions mediate. Indeed, the finding that lateral area 47/12o matures later than medial area 14 supports recently observed developmental differences in the influence of two decision strategies. As introduced above, studies in adult humans and macaques demonstrated that area 47/12o, assigns expected values contingently to choices (Rudebeck *et al.*, 2013b). By contrast, area 14, plays a critical role in expected value coding (Noonan *et al.*, 2011) and biasing choice on the basis of value (Noonan *et al.*, 2010). We recently translated and adapted the multi-option probabilistic decision-making paradigm used in these previous studies and tested a large online sample of adolescents and young adults between 11-35yrs (Wittmann *et al.*, 2021). As the OFC is notably one of the last prefrontal regions to mature across adolescence, we predicted developmental delays in these two behavioural mechanisms. This prediction was borne out for learning strategies supported by the lateral region but not for decision-making strategies supported by medial OFC. Specifically, we found evidence that learning the causal relationships between choice and reward, mediated by the lateral area 47/12o, correlates with age. By contrast, we found no evidence that the children’s choices were influenced by high values of irrelevant alternatives, as mediated by medial OFC.

Understanding OFC grey matter changes across adolescence at this fine-scale is critical to understanding the development of reward guided behaviours more generally, as well as predicting problematic behavioural and mental health outcomes. Broadly, the adolescent OFC shows heightened activity to large rewards and reward predicting cues, compared to adults (Galvan *et al.*, 2006). This enhanced activity to large rewards, particularly in socially rewarding contexts, is associated with reduced cognitive control (Breiner *et al.*, 2018). While too much OFC activity can translate to poor performance, so too can too little OFC grey matter. Diminished cortical thickness in this region (particularly lateral OFC) is associated with adolescent impulsive choice as measured by greater delay discounting (Pehlivanova *et al.*, 2018) and Behavioural Activation and Inhibition questionnaire score (Ide *et al.*, 2020). Collectively, these findings and others have shifted the emphasis of developmental models from broad imbalances between the PFC and subcortical structures to theories suggesting regional neurochemical, structural and functional brain changes within development lead to imbalances within brain circuitry. It is therefore necessary to underlie these theories with detailed examination of brain development with a precise anatomical resolution and to

Altered OFC structure and function in adolescents is associated with a range of mental health disorders. For example, accelerated OFC thinning (particularly more lateral areas) is observed in adolescents who go on to develop depressive symptoms (Bos *et al.*, 2018) and in individuals with bipolar with a history of suicide attempts compared to controls (Huber *et al.*, 2019). While OFC surface morphology differentiates individuals with, or at risk of schizophrenia (Nakamura *et al.*, 2007; Nakamura *et al.*, 2019). Disordered OFC activity is also a common element in mental health conditions, with enhanced activity in tasks in which adolescents make social (Smith *et al.*, 2018) or reward-based decisions. For example, Tegelbeckers and colleagues (2018) observed enhanced signal to large vs small expected rewards in the OFC of ADHD individuals relative to typically developing children/adolescents, with stronger reward-related activity correlated with individual differences in hyperactive/impulsive symptoms in the ADHD group. While the breadth of mental health disorders that the OFC is associated with could indicate a shared underlying dysfunction across all conditions the current apparent overlap and commonalities could also stem from a failure in the precision of which the anatomical foci are reported. If the development of the OFC is not uniform, this heterogeneity has important implications for theories and potential treatment strategies born out of this body of literature. Improving our localisation of these structural and functional differences, and linking them to their unique developmental trajectories, would then facilitate direct comparison to animal models which are already identifying fundamental dissociations in depression and anxiety in the neighbouring ACC’s subregions (Wallis *et al.*, 2017). This is also a crucial step in investigating pharmacological interventions, which is particularly important when there are such few viable treatment options for adolescents (Cipriani *et al.*, 2016).

## Acknowledgements

This work was supported by a grant from the Academy of Medical Sciences. Data were provided by the Human Connectome Project: Young Adult and Development, WU-Minn Consortium funded by the 16 NIH Institutes and Centers that support the NIH Blueprint for Neuroscience Research; and by the McDonnell Center for Systems Neuroscience at Washington University. Finally, we thank Rogier Mars for helpful discussion and comments on the manuscript.

